# Netrin signaling shapes sexually dimorphic neurite branching of a sex-shared interneuron in *C. elegans*

**DOI:** 10.64898/2026.09.29.755526

**Authors:** Da-Hyun Kang, Dongyoung Kim, Geun Park, Byunghyuk Kim

## Abstract

Sexually dimorphic behaviors arise from underlying differences in neural circuit architecture between the sexes, yet the molecular mechanisms that establish these structural differences remain poorly understood. In *Caenorhabditis elegans*, the sex-shared interneuron AVG exhibits sexually dimorphic synaptic connectivity, offering a model for investigating the development of sexually dimorphic neuronal morphology. Here, we identified that AVG displays male-specific neurite branching and demonstrated that the conserved axon guidance cue UNC-6/Netrin and its receptor UNC-40/DCC are essential for this structural remodeling. Cell-specific manipulation of sexual identity revealed that male identity confers branching competence to AVG. Loss of UNC-6 or UNC-40 abolished branch formation, whereas loss of the alternate Netrin receptor UNC-5/Unc5 had no detectable effect. Cell-specific rescue experiments showed that UNC-40 functions in AVG through its UNC-6 binding domains. Furthermore, genetic ablation of the male-specific presynaptic neuron PCA, which expresses *unc-6*, diminished AVG branch length, establishing PCA as a candidate local source of UNC-6. Together, our findings support a model in which presynaptic UNC-6/Netrin cues from male-specific neurons promote dimorphic neurite branching in sex-shared neurons, illustrating how evolutionarily conserved signaling molecules are redeployed during sexual maturation to sculpt sex-specific neural circuits.

## Introduction

Sexual dimorphism manifests across a wide range of biological domains, including physiology, reproductive behaviors, and decision-making. These differences arise from sex-specific gene expression programs and hormonal signals that drive divergent developmental trajectories in each sex (Barr et al., 2018; Knoedler and Shah, 2018; Portman, 2007; Shah et al., 2004). A fundamental outcome of this regulation is the establishment of sexually dimorphic neural circuits, which provide the structural basis for sex-specific behaviors in both invertebrates and vertebrates (Berg et al., 2026; Cachero et al., 2010; Goodwin and Hobert, 2021; Oren-Suissa and Shirangi, 2025; Yang and Shah, 2014). Although anatomical and functional dimorphisms in such circuits are increasingly well-characterized, the molecular mechanisms that generate them—particularly at the level of neurite branching and synapse formation in sex-shared neurons present in both sexes—remain incompletely understood.

In the nematode *Caenorhabditis elegans*, several sex-shared neurons, including DVB, PDB, PHC, PVD, and PVP, undergo striking sex-specific remodeling of neurite branching, synaptic connectivity, and circuit integration during sexual maturation (Hart and Hobert, 2018; Iosilevskii et al., 2025; Kim et al., 2025; Pereira et al., 2019; Serrano-Saiz et al., 2017; Yang et al., 2025). The availability of whole-animal connectomes for both sexes (Cook et al., 2019), combined with high-resolution imaging and powerful genetic tools, makes *C. elegans* a uniquely tractable system for dissecting the cellular and molecular logic of sexually dimorphic circuit assembly at single-neuron resolution.

A recurring principle across animal models is that sex-shared neurons adopt sexually dimorphic morphologies and connectivity patterns under the influence of internal genetic regulators and external inductive cues (Hart and Hobert, 2018; Ito et al., 2016; Kim and Kim, 2022; Kim et al., 2025; Portman, 2017; Sato et al., 2019; Weinberg et al., 2018; Yang and Shah, 2016). Among these conserved pathways, the UNC-6/Netrin signaling system provides a candidate molecular mechanism for shaping sex-specific neural architecture. Comprising the secreted cue UNC-6 and its receptors UNC-40/DCC (Deleted in Colorectal Cancer) and UNC-5/Unc5, the Netrin system plays central roles in axon guidance, neurite outgrowth, and synaptic patterning (Colón-Ramos et al., 2007; Hedgecock et al., 1990; Teichmann and Shen, 2011; Wadsworth, 2002; de Wit and Ghosh, 2016). Beyond its canonical functions in early developmental pathfinding and circuit formation, recent work has implicated UNC-6/Netrin signaling in shaping sexually dimorphic features of sex-shared neurons during sexual maturation. For instance, in the PHB→AVG circuit, male-specific *unc-6* expression maintains synaptic connectivity in males (Weinberg et al., 2018). In the PDB motor neuron, UNC-6 produced by male-specific presynaptic neurons induces elaborate male-specific neurite branching via UNC-40 (Kim et al., 2025). These findings raise the possibility that Netrin-mediated signaling is a broadly utilized strategy to impose sexually dimorphic features onto shared neural frameworks.

Here, we investigate whether UNC-6/Netrin signaling contributes to sexually dimorphic branching in the sex-shared AVG interneuron. AVG has emerged as a key hub in male-specific circuits, exhibiting distinct connectivity that supports tail mechanosensation, locomotion, and mating behavior (Oren-Suissa et al., 2016; Setty et al., 2022). In this study, we characterize the developmental emergence of male-specific AVG neurite branches, examine the contribution of AVG sexual identity, and test the requirement for UNC-6 and its canonical receptors. Furthermore, we identify neighboring male-specific PCA neurons as a candidate local source of UNC-6 and assess the potential behavioral consequences of aberrant male-specific AVG branching. Our findings extend the role of UNC-6/UNC-40 signaling in sexually dimorphic neurite branching to the AVG neuron, providing insights into how conserved guidance cues are redeployed during sexual maturation to sculpt sex-specific structural features and functionalize shared neurons.

## Results

### AVG exhibits male-specific neurite branches

The AVG interneuron is one of the early pioneers of the *C. elegans* ventral nerve cord (VNC), with its cell body located in the retrovesicular ganglion and its axon extending posteriorly along the right VNC into the tailspike (Wadsworth et al., 1996). Electron microscopy (EM) reconstructions indicate that AVG forms neurite branches in the male tail region that are absent in the corresponding hermaphrodite reconstruction (Figure 1A; Cook et al., 2019). Comparison of connectomes between the two sexes also reveals that AVG is embedded in distinct circuit architectures in each sex, receiving numerous synaptic inputs from male-specific neurons in males (Figure 1B). Several of these inputs, including those from PCA and HOA, occur near the male branching region (Figure S1). These anatomical observations motivated an examination of AVG branching by fluorescence imaging.

**Figure 1.**
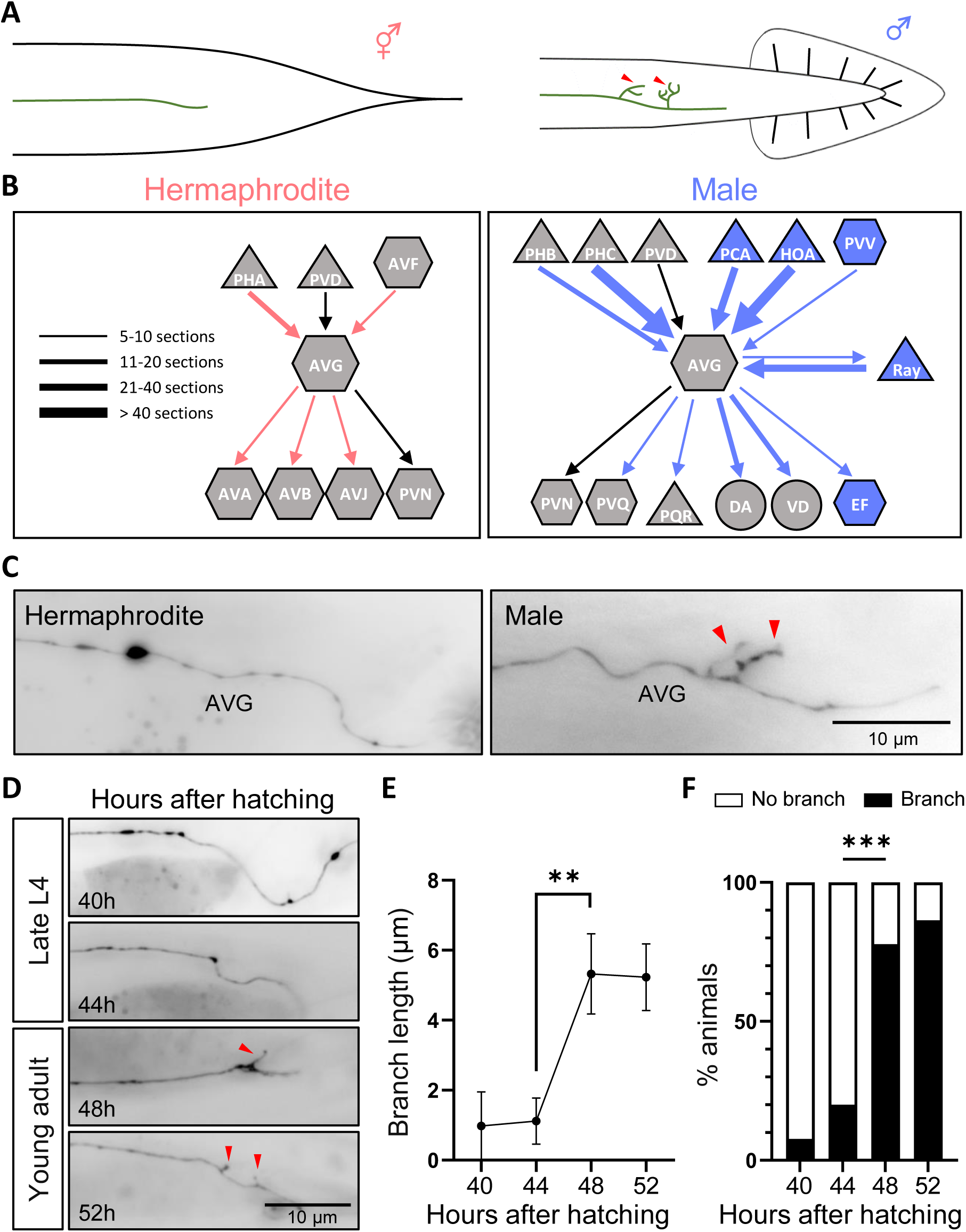
Sexually dimorphic neurite branching of AVG. (A) Schematic representation of the sexually dimorphic morphology of the AVG neuron. Arrowheads indicate male-specific neurite branches. (B) Schematic diagram of AVG synaptic connectivity, based on the number of serial electron micrograph sections. Only chemical synapses present in more than five sections are shown. Sex-shared neurons are depicted in gray, and male-specific neurons in blue. Sex-specific inputs and outputs are indicated by colored arrows. Hexagon, interneuron; triangle, sensory neuron; circle, motor neuron. (C) AVG neurite morphology in adult hermaphrodites and males. The AVG neuron was visualized using the integrated *inx-18p::bfp* reporter (*bxIs29*). Arrowheads indicate male-specific neurite branches. (D) AVG neurite morphology in males at 40, 44, 48, and 52 h after hatching. (E) Quantification of AVG branch length in males at the indicated developmental time points. Error bars are S.E.M. Significance was calculated using the Kruskal-Wallis test with Dunn’s post hoc analysis, \*\**p* < 0.01. *n* ≥ 13. (F) Quantification of the proportion of males with neurite branches at the indicated developmental time points. Significance was calculated using the Chi-square test, \*\*\**p* < 0.001. *n* ≥ 13.

We visualized AVG neurites using a promoter fragment of *inx-18* (*inx-18p*), which drives specific expression in AVG (Oren-Suissa et al., 2016; Wenick and Hobert, 2004). Adult males exhibited one or two short lateral branches near the point at which the right VNC tract diverges in the preanal ganglion, whereas adult hermaphrodites lacked these branches (Figure 1C). The fluorescence observations were consistent with the sex difference apparent in the EM reconstructions (Cook et al., 2019).

### AVG branch morphology diverges between the sexes during sexual maturation

To determine when male AVG branches are formed, we examined AVG morphology at defined stages in synchronized worms. Using 4-hour intervals, we observed that male branches emerge between 44 and 48 hours after hatching, corresponding to the transition from the late L4 stage to adulthood (Figures 1D-1F; Kim et al., 2016). This suggests that male-specific AVG branching is part of the sexually dimorphic remodeling that occurs during sexual maturation.

We also examined AVG morphology in hermaphrodites during this developmental transition. Interestingly, approximately 50% of L4 hermaphrodites exhibited a branch-like protrusion in the region where adult males form branches, whereas adult hermaphrodites lacked this structure (Figure S2). These observations reveal distinct, sex-specific developmental trajectories of AVG morphology during sexual maturation, but do not establish whether the transient hermaphrodite protrusion and the adult male branch share a common developmental origin.

### The sexual identity of AVG influences neurite branching

To elucidate the role of sexual identity in AVG branching, we manipulated the activity of TRA-1, the terminal regulator of the sex determination pathway (Hodgkin, 1987; Zarkower and Hodgkin, 1992). TRA-1 promotes hermaphrodite fate by repressing male-specific gene expression, whereas its degradation, mediated by FEM proteins such as FEM-3, permits male differentiation (Hunter and Wood, 1990; Schvarzstein and Spence, 2006).

We first feminized AVG in males by expressing the intracellular domain of TRA-2 (*tra-2ic*), which antagonizes FEM-3 and thereby stabilizes TRA-1 activity, under the AVG-specific *inx-18* promoter (Mehra et al., 1999; Starostina et al., 2007). In *tra-2ic*-expressing males, both the average branch length and the proportion of animals with branches were reduced, indicating that the male identity of AVG contributes to normal branching (Figures 2A-2C). Next, we masculinized AVG in hermaphrodites by overexpressing *fem-3*. These animals exhibited ectopic neurite outgrowths, although their morphology differed from that observed in wild-type males: whereas wild-type males extend short lateral branches, masculinized hermaphrodite AVG neurons often displayed an axon-like process extending along the left VNC tract (Figures 2A and 2C). Thus, masculinization promotes outgrowth in AVG but does not reproduce normal male branch morphology in a hermaphrodite environment. This result suggests that AVG branch outgrowth is regulated not only by intrinsic sexual identity, but also by the surrounding cellular environment.

**Figure 2.**
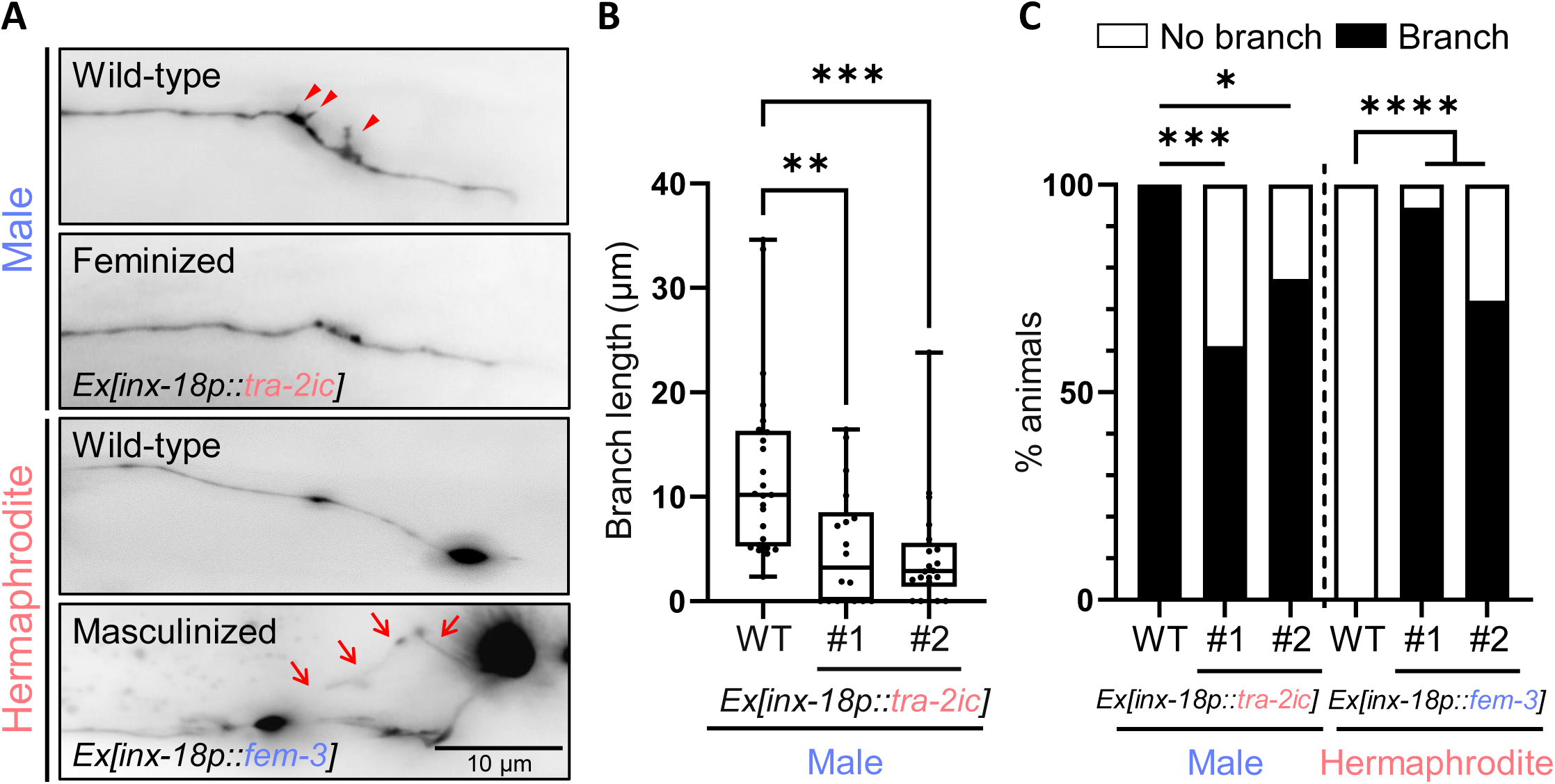
Sexual identity regulates AVG neurite branching. (A) Neurite branch morphology of wild-type or sexually reversed AVG. Arrowheads indicate male-specific AVG neurite branches in wild-type males. Arrows indicate the ectopic, axon-like process in masculinized hermaphrodites. (B) Quantification of AVG neurite branch length in wild-type males and males with feminized AVG. Two independent lines were used for observation. The median is indicated by a horizontal line, the 25th to 75th percentiles are represented by a box, and the minimum and maximum are indicated by whiskers. Significance was calculated using the Kruskal-Wallis test with Dunn’s post hoc analysis. \*\**p* < 0.01; *** *p* < 0.001. *n* ≥ 18. (C) Quantification of the proportion of animals with neurite branches. Two independent lines were used for observation. Significance was calculated using the Chi-square test, * *p* < 0.05; *** *p* < 0.001; **** *p* < 0.0001. *n* ≥ 18.

### Male-specific AVG neurite branching is mediated by UNC-6/Netrin signaling

We previously showed that male-specific branching of the PDB motor neuron is regulated by UNC-6/Netrin signaling (Kim et al., 2025). To test whether a similar molecular logic governs AVG branching, we examined mutant males for UNC-6 and its receptors UNC-40/DCC and UNC-5/Unc5. UNC-6 typically functions through the UNC-40 receptor to attract growing neurites toward the UNC-6 source, whereas UNC-5 mediates repulsive responses (Figure 3A; Hedgecock et al., 1990; Teichmann and Shen, 2011; Wadsworth, 2002).

**Figure 3.**
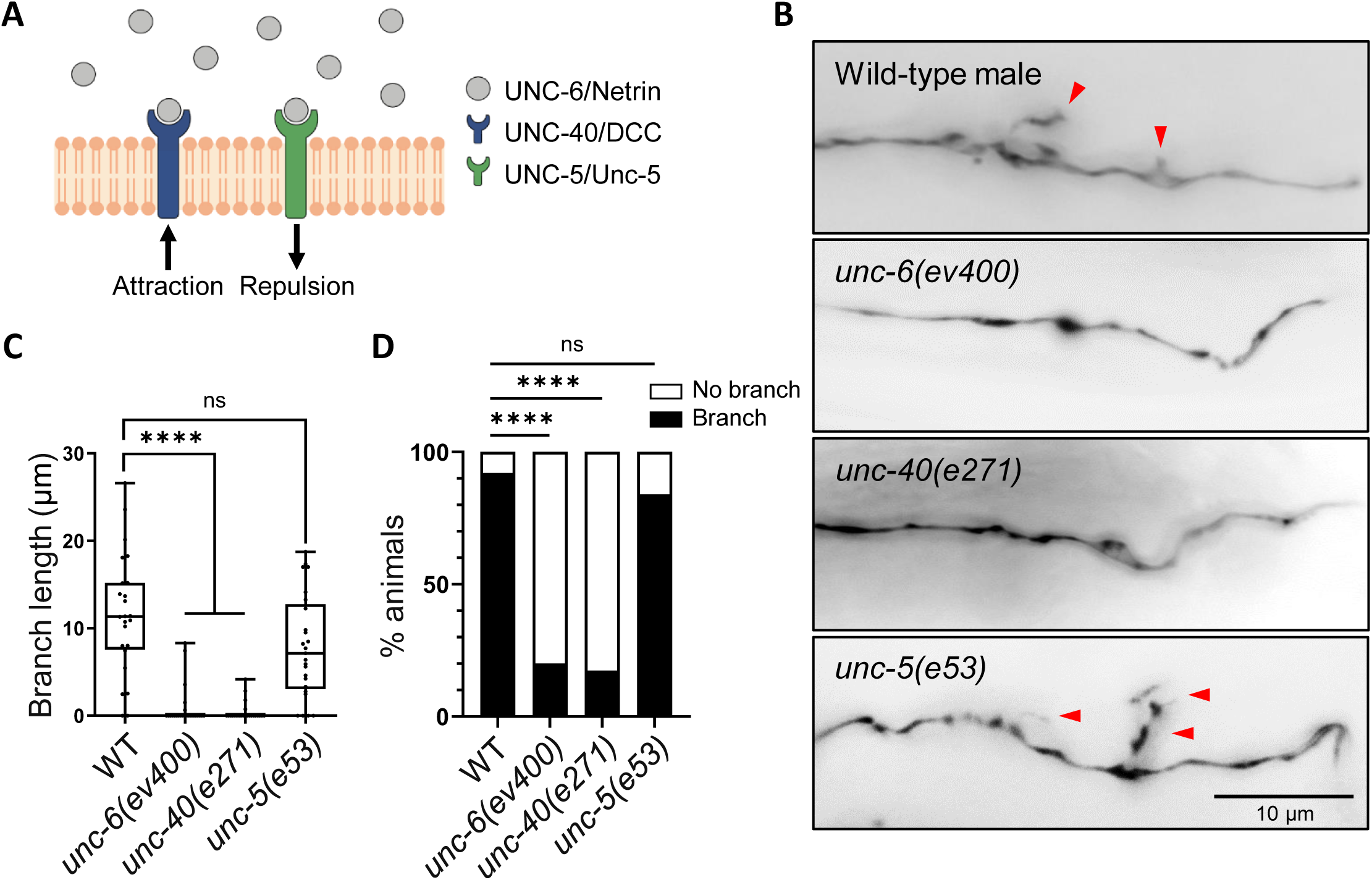
UNC-6/Netrin and its receptor UNC-40/DCC are required for normal male AVG neurite branching. (A) Schematic diagram depicting canonical attractive and repulsive guidance responses mediated by UNC-6/Netrin and its receptors UNC-40/DCC and UNC-5/Unc5. (B) Visualization of male AVG neurons in wild-type and mutants for *unc-6(ev400)*, *unc-40(e271)*, and *unc-5(e53)*. Arrowheads indicate male-specific AVG neurite branches. (C and D) Quantification of AVG branch length (C) and the proportion of animals with neurite branches (D) in mutant strains. Significance was calculated using the Kruskal-Wallis test with Dunn’s post hoc analysis (C) or the Chi-square test (D), \*\*\*\**p* < 0.0001; ns, not significant. *n* ≥ 23.

In *unc-6(ev400)* mutants, AVG branches were dramatically reduced in both length and frequency, indicating that UNC-6 is required for branch formation (Figures 3B-3D). A similar phenotype was observed in *unc-40(e271)* mutants, in which AVG branches were rarely detectable and were significantly shortened when present (Figure 3B-3D). By contrast, *unc-5(e53)* mutants exhibited normal AVG branches indistinguishable from those of wild-type controls (Figure 3B-3D). Together, these results demonstrate that UNC-6/UNC-40 signaling, but not UNC-5, is required for normal male AVG branching.

### UNC-40 acts in AVG and requires UNC-6 binding domains to promote branching

To determine whether UNC-40 functions cell-autonomously in AVG, we performed a rescue experiment in *unc-40* mutants. Expression of *unc-40* specifically in AVG (*inx-18p::unc-40*) restored AVG branching, including both branch length and the proportion of animals with branches, supporting that UNC-40 acts in AVG to promote neurite branch formation (Figure 4). To test whether this function depends on UNC-6 binding, we expressed a truncated *unc-40* cDNA lacking the FNIII4-5 domains, which are essential for its interaction with UNC-6 (Meijers et al., 2020). This construct failed to rescue the mutant phenotype, indicating that UNC-40 promotes AVG branching through its UNC-6 binding domains (Figure 4). This cell-autonomous role is consistent with prior studies showing that UNC-40 interacts with UNC-6 to guide neurite outgrowth and axonal pathfinding in other *C. elegans* neurons (Chan et al., 1996; Kim et al., 2025; Teichmann and Shen, 2011).

**Figure 4.**
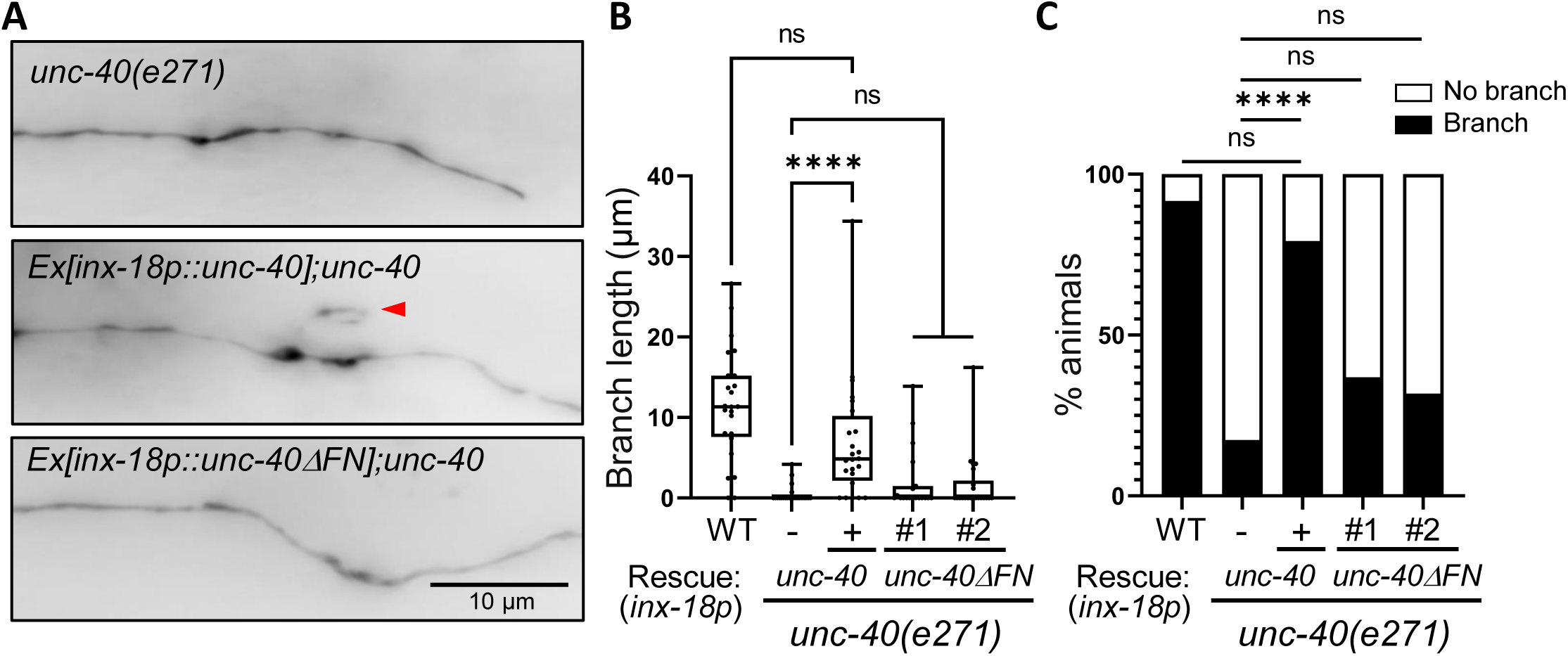
UNC-40 functions in AVG and requires UNC-6 binding domains. (A) Visualization of male AVG neurons in *unc-40(e271)* mutants and transgenic *unc-40(e271)* animals expressing either full-length *unc-40* cDNA (*inx-18p::unc-40*) or a truncated *unc-40* cDNA lacking the FNIII4-5 domains (*inx-18p::unc-40ΔFN*) specifically in AVG. Arrowheads indicate male-specific AVG neurite branches. (B and C) Quantification of AVG branch length (B) and the proportion of animals with neurite branches (C) in rescue lines. Two independent lines were used to observe the rescue phenotype of *inx-18p::unc-40ΔFN*. Quantification data for wild-type and *unc-40* mutants are as in Figure 3 for comparison. Significance was calculated using the Kruskal-Wallis test with Dunn’s post hoc analysis (B) or the Chi-square test (C), \*\*\*\**p* < 0.0001; ns, not significant. *n* ≥ 19.

### The male-specific PCA neuron is a candidate local source of UNC-6

Having established that UNC-6 is required for male-specific AVG neurite branching, we next investigated potential cellular sources of UNC-6 located near AVG branches. Connectome data identified the male-specific neurons PCA and HOA and the sex-shared neuron PHC as major presynaptic partners near the branching region (Figure S1). EM reconstructions from WormWiring (https://wormwiring.org/) showed spatial overlap between these neurites and AVG branches in this region (Figure 5A; Cook et al., 2019; Jarrell et al., 2012). Fluorescence imaging further confirmed close apposition of these neurites (Figures 5B).

**Figure 5.**
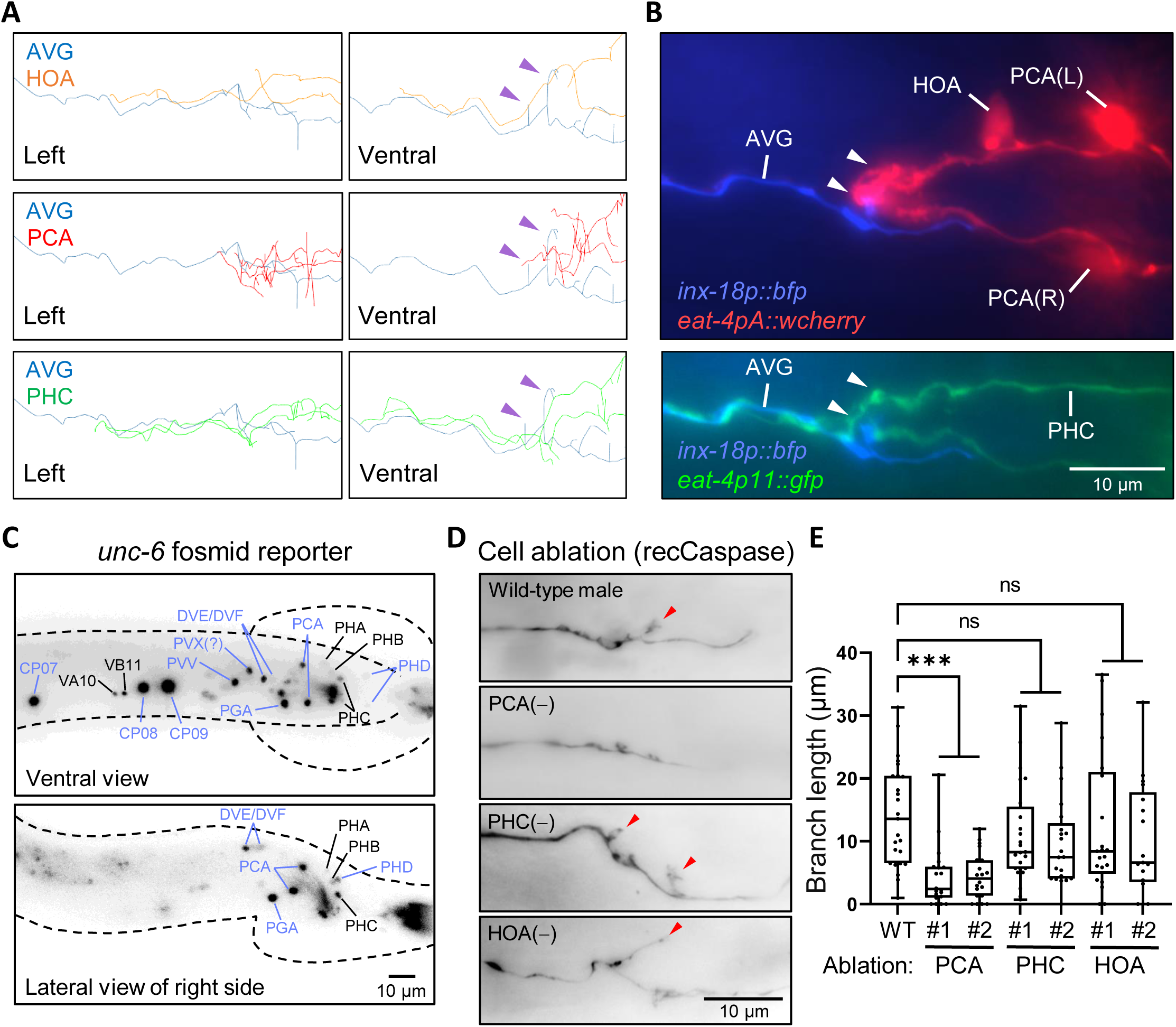
PCA neurons express *unc-6* and are required for AVG neurite branching. (A) Skeleton diagrams of male-specific AVG neurite branches (blue) with presynaptic partners HOA (orange), PCA (red), and PHC (green), based on electron micrograph reconstructions from WormWiring (http://wormwiring.org/). Ventral views show areas with significant neurite overlap (arrowheads). (B) Representative fluorescence images of male-specific AVG neurite branches (*inx-18p::bfp*, blue) showing neurite overlap with HOA and PCA (*eat-4pA::wcherry*, red) and PHC (*eat-4p11::gfp*, green). Arrowheads indicate the sites of neurite overlap. (C) Visualization of *unc-6* expression in the adult male tail using a fosmid-based transcriptional reporter (*otIs638[unc-6::SL2::NLS::YFP::H2B]*). Male-specific neurons (blue) and sex-shared neurons (black) are indicated. Note that *unc-6* expression was observed in PCA and PHC, but not in HOA. (D) Fluorescence images of AVG neurites in wild-type males and males following genetic ablation of PCA, PHC, or HOA. Cell-specific ablation was conducted for PCA with two split-caspase constructs (*eat-4pA::tu813* and *syg-1p::tu814*), for PHC (*eat-4p11::tu813* and *eat-4p11::tu814*), or for HOA (*eat-4pA::tu813* and *sax-3p::tu814*). (E) Quantification of AVG branch length in cell-ablated animals. Two independent lines were used for observation. Significance was calculated using the Kruskal-Wallis test with Dunn’s post hoc analysis. \*\*\**p* < 0.001; ns, not significant. *n* ≥ 17.

To identify which presynaptic neurons express *unc-6*, we examined a fosmid-based *unc-6* transcriptional reporter (*unc-6::SL2::NLS::YFP::H2B*) (Weinberg et al., 2018). This reporter was detected in PCA and PHC but not in HOA (Figure 5C). Although the fosmid reporter did not show detectable signal in HOA, recent single-cell RNA sequencing datasets suggest potential *unc-6* expression in HOA as well (Haque et al., 2025; Morillo et al., 2025).

To test the functional requirement of the three presynaptic neurons, we genetically ablated each neuron using split-caspase constructs (Figure S3; Chelur and Chalfie, 2007). Ablation of the male-specific PCA neurons significantly reduced AVG branch length, whereas ablation of HOA or PHC had no detectable effect on branch length (Figures 5D and 5E). Together, these results suggest that PCA is a major local presynaptic contributor to UNC-6-dependent AVG branching, although we cannot rule out the possibility that additional sources of UNC-6 or other signals also contribute to branch formation.

### Aberrant AVG neurite branches correlate with defects in male behavior

Finally, we asked whether male-specific AVG neurite branches contribute to male behaviors. Previous studies showed that ablation or silencing of the male AVG neuron impairs tail mechanosensation, locomotion, and vulva location during mating (Oren-Suissa et al., 2016; Setty et al., 2022). To dissect the functional significance of the branches, we analyzed males with feminized AVG, which exhibit reduced neurite branching without eliminating the AVG cell body (Figure 2). Males with feminized AVG showed a marked reduction in vulva location efficiency during mating (Figure 6A). In addition, these males displayed a reduced tail-touch response and decreased locomotion speed, whereas their hermaphrodite counterparts showed no significant change in either measure (Figures 6B and 6C). These findings suggest that male-specific AVG neurite branches may contribute to the proper execution of male behaviors, likely through the male-specific synaptic connections they support.

**Figure 6.**
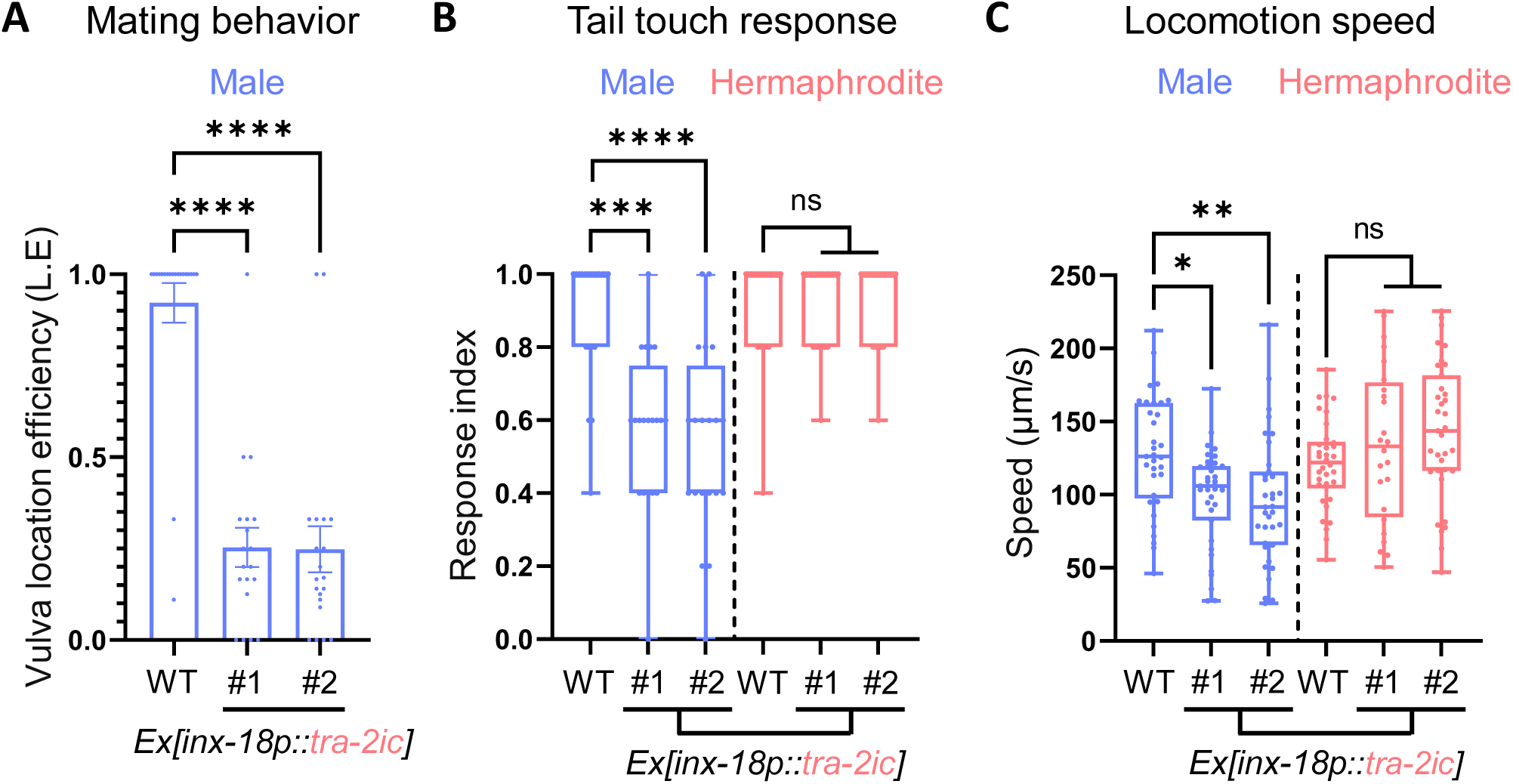
Aberrant AVG neurite branches correlate with defects in male behavior. (A) Vulva location efficiency of wild-type males and males with cell-specifically feminized AVG (*inx-18p::tra-2ic*). Two independent lines were used for observation. Error bars are S.E.M. Each dot represents individual animal. Significance was calculated using the Kruskal-Wallis test with Dunn’s post hoc analysis. \*\*\*\**p* < 0.0001. *n* ≥ 19. (B and C) Tail touch response index (B) and locomotion speed (C) in wild-type and AVG-feminized animals. AVG-feminized hermaphrodites were used as controls for potential effects of the feminization on behavior. Significance was calculated using the Kruskal-Wallis test with Dunn’s post hoc analysis. \**p* < 0.05; \*\**p* < 0.01; \*\*\**p* < 0.001; \*\*\*\**p* < 0.0001. *n* ≥ 20 in (B) and *n* ≥ 24 in (C).

## Discussion

Sexually dimorphic neural circuits enable sex-specific behaviors across diverse organisms, yet the molecular mechanisms sculpting their distinct architectures remain only partially understood. In this study, we demonstrate that UNC-6/Netrin and UNC-40/DCC signaling establishes sexually dimorphic neurite branching in the sex-shared AVG interneuron. Our findings identify that AVG sexual identity influences neurite outgrowth and that the male-specific presynaptic neuron PCA contributes to normal branching. Together, these data support a model in which an intrinsic sex-dependent program and local extracellular cues contribute to AVG neurite remodeling. In this model, local UNC-6 supplied by the male-specific PCA neuron acts through UNC-40 in AVG to promote lateral branch formation, thereby functionalizing AVG within male-specific circuits (Figure S4).

The AVG neuron extends short lateral branches in adult males but not adult hermaphrodites. This structural difference accompanies sexually dimorphic synaptic connectivity, consistent with the broader association between sex-specific neuritogenesis and circuit configuration (Cook et al., 2019; Hart and Hobert, 2018; Kim et al., 2025; Serrano-Saiz et al., 2017). Developmental profiling shows that male-specific AVG branches emerge between 44 and 48 hours after hatching, around the late L4-to-adult transition. This timing aligns with the assembly period of male-specific circuits during sexual maturation (Emmons, 2014; Sulston et al., 1980). In contrast, branch-like protrusions were observed in approximately half of L4 hermaphrodites, but were absent in adults. These observations suggest that adult AVG dimorphism may involve not only the emergence of male branches but also the loss or remodeling of transient structures in hermaphrodites. The transient hermaphrodite protrusion could represent a morphological precursor associated with previously described sex-specific synaptic pruning of PHB→AVG synapses in hermaphrodites (Oren-Suissa et al., 2016). However, its relationship to the sex-specific elimination of these synapses remains unresolved.

Manipulation of AVG sexual identity indicates that feminization reduces neurite branching in males, whereas masculinization induces ectopic processes in hermaphrodites. These reciprocal manipulations implicate intrinsic sexual identity in AVG branch outgrowth. Importantly, masculinized hermaphrodite AVG neurons did not acquire the normal male branch morphology; they often extended an axon-like process along the left VNC tract. This distinction suggests that male identity alone is insufficient to specify the spatial pattern of normal male branches. The surrounding neuronal environment may influence branch position or trajectory, although the relevant extracellular determinants have not been resolved. The relationship between sexual identity and UNC-6/UNC-40 signaling also remains open: male identity could alter receptor abundance, localization, or downstream signaling. Our experiments establish contributions from both sexual identity and Netrin signaling without defining their mechanistic relationship.

Our genetic and cell-specific rescue experiments support a cell-autonomous role of UNC-40 in AVG branching that relies on its Netrin-binding domains. Cell ablation and expression analyses further implicate the male-specific sensory neuron PCA as a candidate presynaptic source of UNC-6. This parallel with the sex-shared PDB motor neuron, in which male-specific CP neurons supply UNC-6 to induce UNC-40-dependent PDB branching (Kim et al., 2025), highlights a broader principle: the conserved Netrin signaling is repeatedly redeployed during sexual maturation to sculpt male-specific neurite branches in sex-shared neurons, with male-specific neurons serving as local sources.

Behavioral characterization of males with feminized AVG reveals defects in vulva location, tail-touch responsiveness, and locomotion speed, consistent with previously described phenotypes of AVG ablation or silencing (Oren-Suissa et al., 2016; Setty et al., 2022). Because feminized AVG retains the cell body, these defects correlate branch structure with circuit function regulating these behaviors. However, as cell-specific feminization is also known to alter transcriptional profiles, synaptic connectivity, and physiological properties (Fagan et al., 2018; Oren-Suissa et al., 2016; Serrano-Saiz et al., 2017), the behavioral phenotypes we observed cannot be attributed exclusively to the loss of the branch. Developing tools to selectively sever or prevent branch formation without altering sexual identity will be an important future step toward establishing direct causality.

In conclusion, our findings show that UNC-6/Netrin not only functions as a global gradient or positional cue during early development, but also appears to be modularly co-opted to shape sex-specific circuits by directing local, sex-specific neurite branching. Gating access to conserved guidance cues via sex-determination pathways could provide a flexible and efficient strategy to diversify neural architecture and behavioral repertoires.

## Methods

### C. elegans strains

All strains were maintained according to standard methods (Brenner, 1974). Wild-type strains used in this study contained *bxIs29[eat-4p11::gfp, eat-4pA::wcherry, inx-18p::bfp]* to visualize AVG, PCA, and PHC neurons (Kim and Emmons, 2017). All strains contained the *him-5(e1490)* allele to obtain high proportions of males. Mutant alleles used were *unc-6(ev400)*, *unc-40(e271),* and *unc-5(e53)*. Worms were grown at 21 °C on nematode growth media (NGM) plates seeded with bacteria (*Escherichia coli* OP50) as a food source. All transgenic strain information is listed in Table S1.

### Molecular cloning and constructs

To generate pBK163 (*inx-18p::tra-2ic::sl2::mCherry*) for AVG feminization, *tra-2ic::sl2::mCherry* was digested from pBK137 (*kal-1p9::tra-2ic::sl2::mCherry*) with the restriction enzymes SphI and XmaI. This fragment was ligated in SphI and XmaI-digested pBK83 (*inx-18p::spYFP::Δrig-6d-4fn3*).

To generate pBK164 (*inx-18p::fem-3::sl2::mCherry*) for AVG masculinization, *fem-3::sl2::mCherry* was PCR-amplified from pBK132 (*srh-142p::fem-3::sl2::mCherry*, a gift from Portman, D. S.) with the addition of the restriction sites of NheI and KpnI in the primers. This PCR product was digested and ligated in NheI and KpnI-digested pBK162 (*inx-18p::unc-40ΔFNIII4-5domain*).

To generate pBK159 (*inx-18p::unc-40cDNA*) for *unc-40* rescue, *unc-40cDNA* was digested from pBK115 (*rol-6Bb::unc-40cDNA*) with the restriction enzymes SphI and XmaI. This fragment was ligated in SphI and XmaI-digested pBK45 (*inx-18p::casy-1*).

To generate pBK162 (*inx-18p::unc-40ΔFNIII4-5domain*) for *unc-40ΔFNIII4-5domain* rescue, *unc-40ΔFNIII4-5domain* was digested from pBK109 (*kal-1p9::unc-40ΔFNIII4-5domain*) with the restriction enzymes SphI and XmaI. This fragment was ligated in SphI and XmaI-digested pBK83 (*inx-18p::spYFP::Δrig-6d-4fn3*).

To generate pBK166 (*eat-4pA::tu813*) for PCA and HOA cell death, *eat-4pA* was PCR-amplified from pBK71 (*eat-4pA::sax-7S*) with the addition of the restriction enzyme sites of SphI and SbfI in the primers. This PCR product was digested and ligated in SphI and SbfI-digested pBK125 (*rol-6Bb::tu813*).

To generate pBK167 (*syg-1p::tu814*) for PCA cell death, *syg-1p* was PCR-amplified from worm lysis with the addition of the restriction sites SphI and SbfI in the primers. This PCR product was digested and ligated in SphI and SbfI-digested pBK143 (*flp-21p::tu814*).

To generate pBK168 (*eat-4p11::tu813*) for PHC cell death, *eat-4p11* was PCR-amplified from pBK72 (*eat-4p11::sax-7S*) with the addition of the restriction sites of SphI and SbfI in the primers. This PCR product was digested and ligated in SphI and SbfI-digested pBK142 (*rol-6Bb::tu813*).

To generate pBK169 (*eat-4p11::tu814*) for PHC cell death, *eat-4p11* was PCR-amplified from pBK72 (*eat-4p11::sax-7S*) with the addition of the restriction sites of SphI and SbfI in the primers. This PCR product was digested and ligated in SphI and SbfI-digested pBK143 (*flp-21p::tu814*).

To generate pBK170 (*sax-3p::tu814*) for HOA cell death, *sax-3p* was PCR-amplified from worm lysis with the addition of the restriction sites SphI and SbfI in the primers. This PCR product was digested and ligated in SphI and SbfI-digested pBK124 (*rol-6Bb::tu814*).

### Synchronized worm preparation

To obtain male populations of defined age, worms were synchronized as previously described (Kim et al., 2016). In brief, gravid hermaphrodites were allowed to lay eggs overnight and then washed off the plate with M9 buffer, together with any larvae, leaving only eggs adhering to the agar surface. Newly hatched L1 worms were harvested at 1-h intervals by gently washing them from the plate with M9 buffer. Harvested worms were grown at 20 °C and imaged at defined time intervals (40, 44, 48, and 52 h post-hatching).

### Microscopy and neurite tracing

Worms were anesthetized in M9 buffer containing 50-100 mM of sodium azide (NaN_3_) and mounted on 5% (w/v) agarose pad in deionized water on glass slides. We used 1-day-old adult males unless otherwise indicated. Worms were imaged by high-resolution fluorescence microscopy (Zeiss Axio Imager.Z2). Images were processed using AxioVision (ver. 4.03) and image reconstruction was performed using Z-projection plug-in in ImageJ version 2 package, FIJI of NIH ImageJ package (ver. 2.13.1). Z-stack images were loaded into the Simple Neurite Tracer (SNT) plug-in in FIJI (Arshadi et al., 2021). The length of AVG neurite branches was measured using SNT.

### Mating behavior assay

L4 males were isolated and kept for 16-24 h until they reached the adult stage. NGM plates were prepared with a 2-μl drop of 10 × concentrated, freshly prepared OP50 *E. coli* culture and up to five 1-day-old adult *unc-31(e169)* hermaphrodites. A single adult male was placed on each plate and its mating behavior was scored for 5 min or until ejaculation, whichever came first. Vulva location efficiency was calculated as 1 / (number of encounters required to achieve a stop), where a successful encounter was defined as a complete stop of the male tail at the vulva for at least 10 s (Peden and Barr, 2005).

### Tail touch assay

Tail mechanosensation was measured as previously described (Setty et al., 2022). L4 males or hermaphrodites were isolated and kept for 16-24 h until they reached adulthood. Single 1-day-old adult worms were transferred onto NGM plates seeded with 30 μl OP50. After 30 min of habituation, animals that were not moving or moving very slowly were touched on the tail with a flattened platinum wire pick, with an interval of at least 10 s between five trials per worm. Each trial was scored 1 if the worm moved forward in response to touch and 0 if it did not; the response index was calculated as the average of the five trials.

### Movement tracking

One-day-old adult males or hermaphrodites were transferred to food-free NGM plates and allowed 5 min to acclimate. Animals were recorded for 30 s using a Leica MC170 HD microscope camera controlled by Leica LAS EZ imaging software (ver. 3.4.0). Locomotory behavior was quantified using Deep-Worm-Tracker, which employs YOLOv5 object detection and StrongSORT multi-object tracking algorithms (Banerjee et al., 2023). Worm centroids were calculated from bounding box coordinates and smoothed using a 5-frame moving average to reduce measurement noise. Frame-to-frame distances were calculated using the Euclidean distance formula, with movements exceeding 30 pixels per frame excluded as outliers. Movement speed was calculated as total distance traveled divided by tracking duration and converted from pixel measurements to physical units using a calibration factor of 7.8 μm/pixel and 30 fps video acquisition.

### Statistical analysis

Statistical analysis was performed using GraphPad Prism software (version 10.0.0.15., RRID:SCR_002798). Specific statistical methods used are described in the figure legends.

## Acknowledgements

We thank Minkyung Shin and Seungwon Oh for their assistance in preparing experimental reagents. Some strains were provided by the CGC, which is funded by NIH Office of Research Infrastructure Programs (P40 OD010440). This work was supported by the National Research Foundation of Korea (NRF) grant funded by the Korean government (MSIT) (RS-2026-25481743).

## Supplementary figure legends

**Figure S1.**
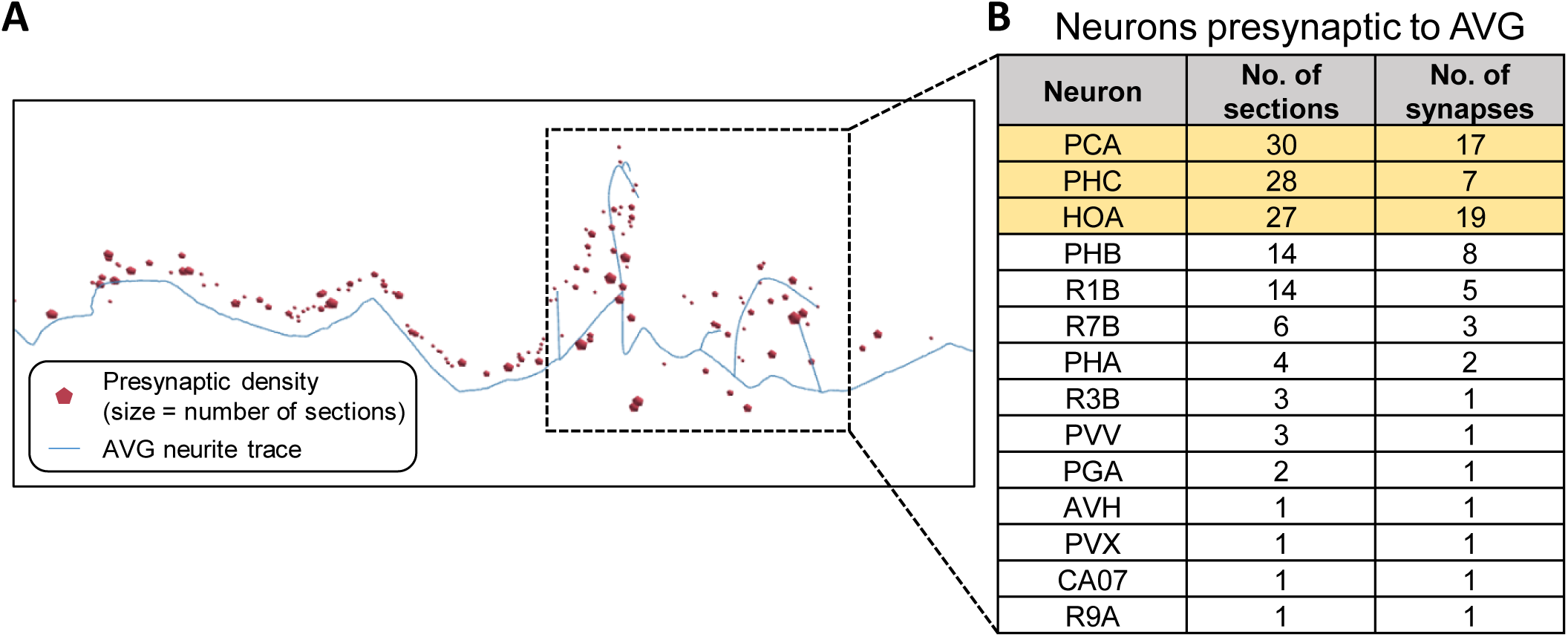
Presynaptic partners of AVG near the male-specific branching region. (A) Skeleton trace of the male AVG axon showing the spatial distribution of presynaptic densities along its neurite. Red dots indicate sites at which presynaptic partners contact AVG; dot size is proportional to the number of serial electron micrograph sections. The dashed box outlines the region containing male-specific neurite AVG branches. (B) Summary table of presynaptic neurons making synapses with AVG within the branching region. The table lists the total number of electron micrograph sections and the number of synapses for each partner.

**Figure S2.**
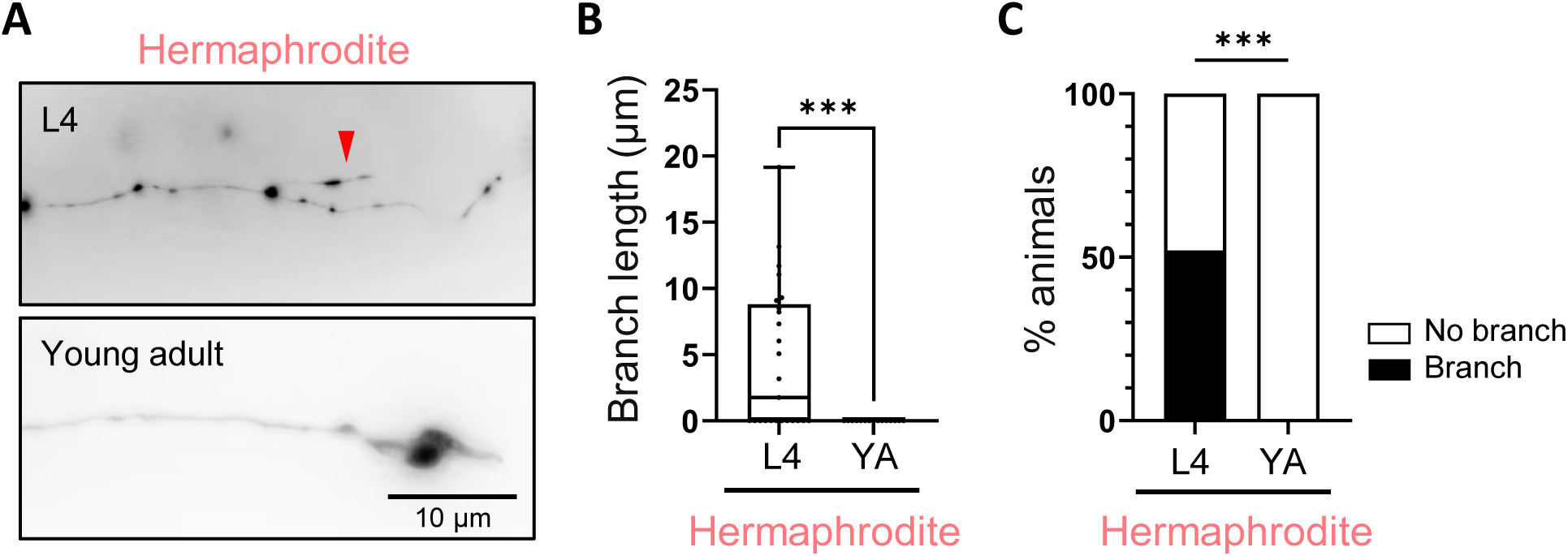
Transient AVG neurite branching during hermaphrodite maturation. (A) Representative morphology of AVG neurites in hermaphrodites at the L4 larval and young adult (YA) stages. The arrowhead marks a transient neurite branch present during the L4 stage. (B and C) Quantification of AVG branch length (B) and the proportion of animals with AVG neurite branches (C) in L4 and YA hermaphrodites. Significance was calculated using the Mann-Whitney test (B) or the Chi-square test (C). \*\*\**p* < 0.001; *n* ≥ 15.

**Figure S3.**
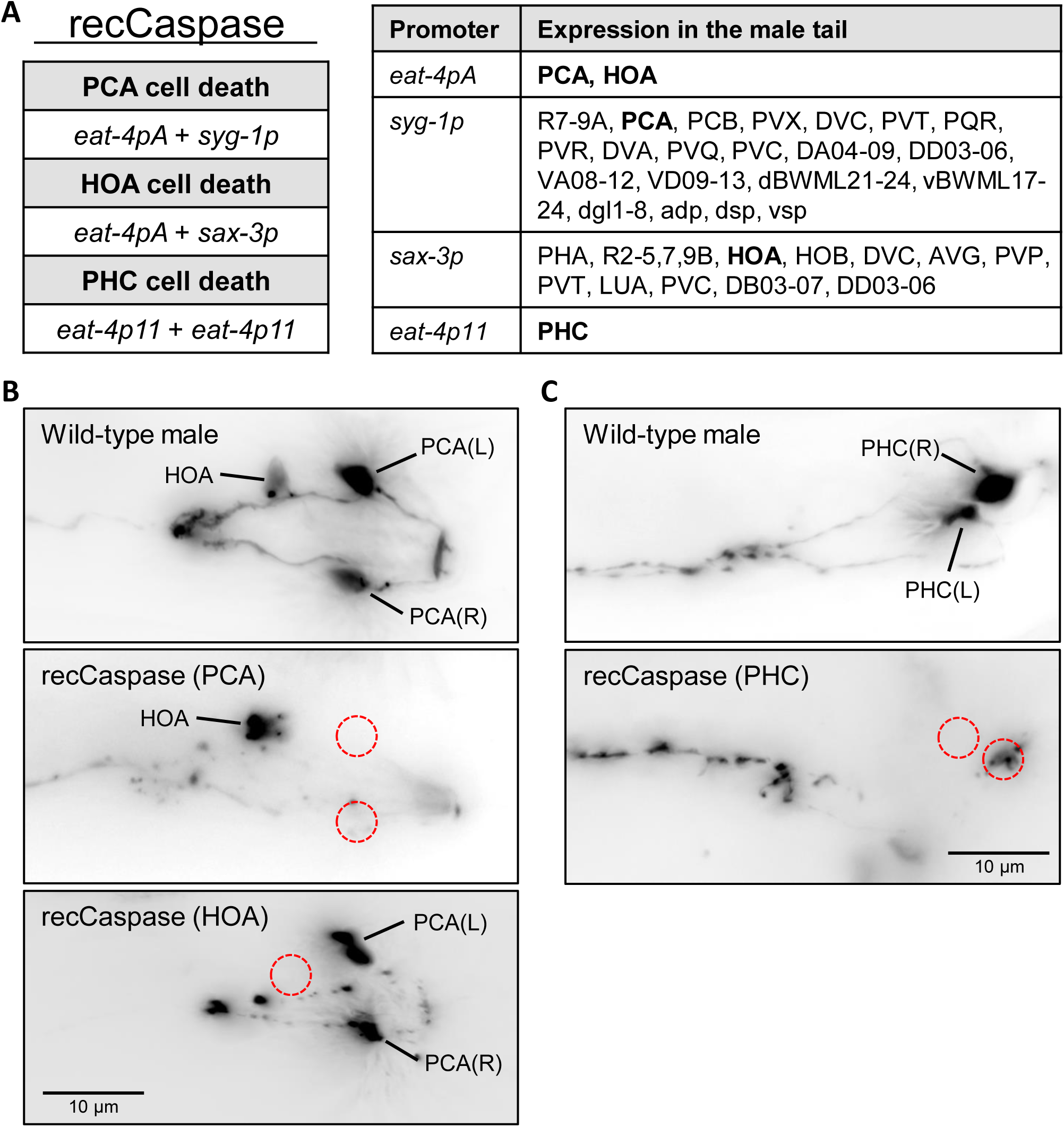
Constructs and validation of cell-specific neuronal ablation. (A) Summary of split-caspase constructs used for cell-specific ablation of PCA, HOA, and PHC neurons. The left table lists the promoter combinations used to drive expression of the two split-caspase components. The right table lists reported cellular expression profiles for each promoter (Kim and Emmons, 2017; Sakai et al., 2023). (B and C) Representative fluorescence images of the adult male tail region for PCA or HOA ablation (B) and PHC ablation (C). Dashed circles outline the expected anatomical positions of the ablated neuronal cell bodies.

**Figure S4.**
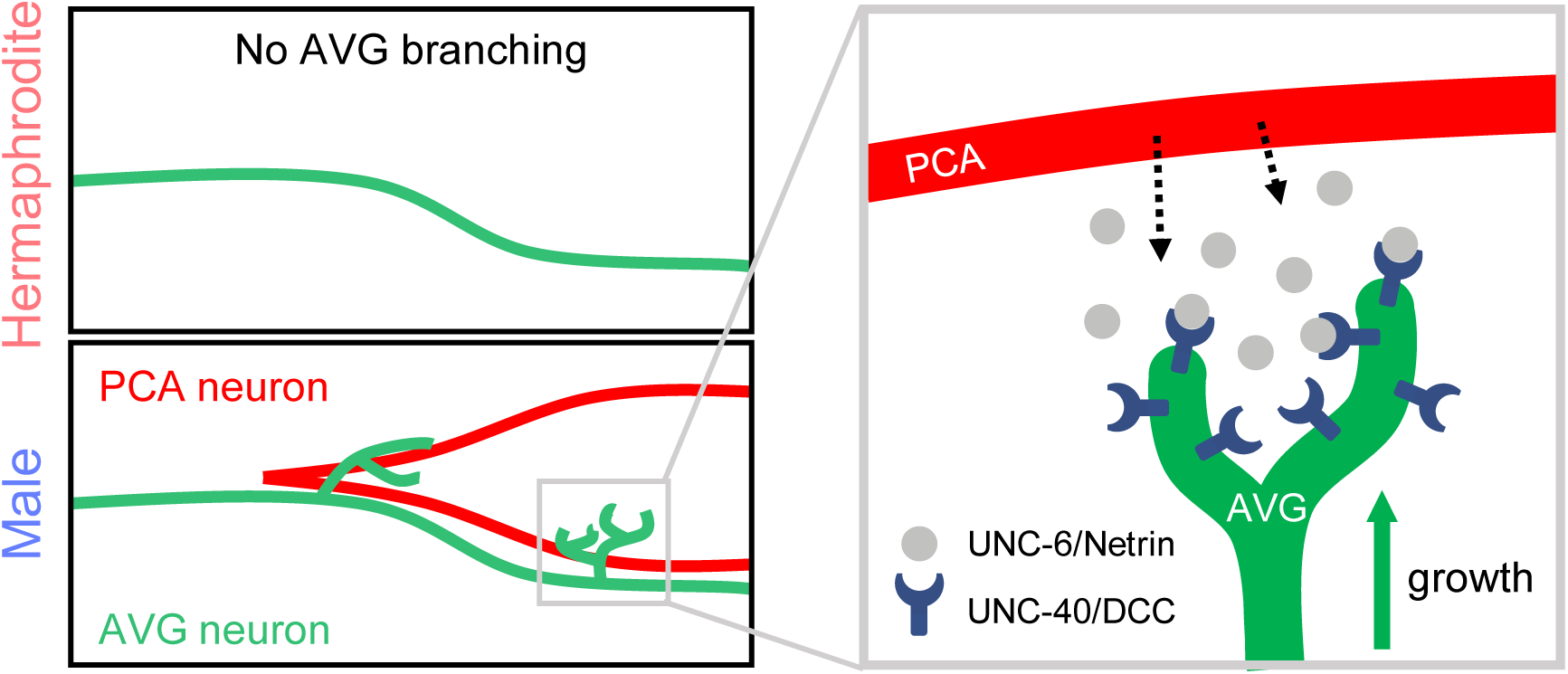
Proposed model for UNC-6/Netrin-mediated regulation of male-specific neurite branching. In hermaphrodites, AVG does not form lateral branches. In males, local presynaptic partners —specifically the male-specific PCA neuron—are positioned adjacent to AVG near the potential branching site. UNC-6/Netrin released from PCA interacts with UNC-40/DCC on the plasma membrane of AVG to promote lateral neurite branching.

**Table S1.** Transgenic strains used in this study.

| Strain name | Transgene name | DNA on array | Strain background | Source |
| --- | --- | --- | --- | --- |
| EM1462 | *bxIs29* | *eat-4p11::gfp + eat-4pA::wcherry + inx-18p::bfp* | *him-5(e1490)V* | Kim and Emmons (2017) |
| BHK186 | *goyEx143; bxIs29* | *inx-18p::tra-2ic::sl2::mcherry (20ng/ul) + ttx-3p::gfp (50ng/ul) + DNA ladder(1kb plus) (30ng/ul) – Line #1* | *him-5(e1490)V* | This study |
| BHK191 | *goyEx143; bxIs29* | *inx-18p::tra-2ic::sl2::mcherry (20ng/ul) + ttx-3p::gfp (50ng/ul) + DNA ladder(1kb plus) (30ng/ul) – Line #2* | *him-5(e1490)V* | This study |
| BHK185 | *goyEx142; bxIs29* | *inx-18p::fem-3::sl2::mcherry (20ng/ul) + ttx-3p::GFP (50ng/ul) + DNA ladder(1kb plus) (30ng/ul) – Line #1* | *him-5(e1490)V* | This study |
| BHK188 | *goyEx142; bxIs29* | *inx-18p::fem-3::sl2::mcherry (20ng/ul) + ttx-3p::GFP (50ng/ul) + DNA ladder(1kb plus) (30ng/ul) – Line #2* | *him-5(e1490)V* | This study |
| BHK151 | *bxIs29* | *eat-4p11::gfp + eat-4pA::wcherry + inx-18p::bfp* | *him-5(e1490)V; unc-40(e271)I* | This study |
| BHK161 | *bxIs29* | *eat-4p11::gfp + eat-4pA::wcherry + inx-18p::bfp* | *him-5(e1490)V; unc-6(ev400)X* | This study |
| BHK179 | *bxIs29* | *eat-4p11::gfp + eat-4pA::wcherry + inx-18p::bfp* | *him-5(e1490)V; unc-5(e53) IV* | This study |
| BHK182 | *goyEx138; bxIs29* | *inx-18p::unc-40cDNA (25ng/ul) + ttx-3p::GFP (25ng/ul) + DNA ladder (50ng/ul)* | *him-5(e1490)V; unc-40(e271)* | This study |
| BHK187 | *goyEx144; bxIs29* | *inx-18p::ΔFNIII domain 4-5 (25ng/ul) + ttx-3p::GFP (25ng/ul) + DNA ladder(1kb plus) (50ng) – Line #1* | *him-5(e1490)V; unc-40(e271)* | This study |
| BHK190 | *goyEx144; bxIs29* | *inx-18p::ΔFNIII domain 4-5 (25ng/ul) + ttx-3p::GFP (25ng/ul) + DNA ladder(1kb plus) (50ng) – Line #2* | *him-5(e1490)V; unc-40(e271)* | This study |
| BHK86 | *otIs638* | *unc-6::SL2::NLS::YFP::H2B + ttx-3p::mcherry + pha-1(+)* | *him-5(e1490)V* | CGC |
| BHK198 | *goyEx146; bxIs29* | *eat-4pA::tu813 (25ng/ul) + syg-1p::tu814 (25ng/ul) + ceh-22p::GFP (25ng/ul) + DNA ladder(1kb plus) (25ng/ul) – Line #1* | *him-5(e1490)V* | This study |
| BHK199 | *goyEx146; bxIs29* | *eat-4pA::tu813 (25ng/ul) + syg-1p::tu814 (25ng/ul) + ceh-22p::GFP (25ng/ul) + DNA ladder(1kb plus) (25ng/ul) – Line #2* | *him-5(e1490)V* | This study |
| BHK201 | *goyEx147; bxIs29* | *eat-4p11::tu813 (25ng/ul) + eat-4p11::tu814 (25ng/ul) + ceh-22p::GFP (25ng/ul) + DNA ladder(1kb plus) (25ng/ul) – Line #1* | *him-5(e1490)V* | This study |
| BHK203 | *goyEx147; bxIs29* | *eat-4p11::tu813 (25ng/ul) + eat-4p11::tu814 (25ng/ul) + ceh-22p::GFP (25ng/ul) + DNA ladder(1kb plus) (25ng/ul) – Line #2* | *him-5(e1490)V* | This study |
| BHK208 | *goyEx150; bxIs29* | *eat-4pA::tu813 (25ng/ul) + sax-3p::tu814 (25ng/ul) + ttx-3::GFP (25ng/ul) + DNA ladder(1kb plus) (25ng/ul) – Line #1* | *him-5(e1490)V* | This study |
| BHK209 | *goyEx150; bxIs29* | *eat-4pA::tu813 (25ng/ul) + sax-3p::tu814 (25ng/ul) + ttx-3::GFP (25ng/ul) + DNA ladder(1kb plus) (25ng/ul) – Line #2* | *him-5(e1490)V* | This study |

**Table S2.** The number of animals used in each experiment.

|  |  |
| --- | --- |
| Figures 1E and 1F | Wild-type 40 hours after hatching $n = 13$, 44 hours $n = 20$, 48 hours $n = 18$, 52 hours $n = 22$ |
| Figure 2B | Wild-type $n = 25$, *Ex[inx-18p::tra-2ic]* #1 $n = 18$, *Ex[inx-18p::tra-2ic]* #2 $n = 22$ |
| Figure 2C | Wild-type male $n = 25$, *Ex[inx-18p::tra-2ic]* #1 $n = 18$, *Ex[inx-18p::tra-2ic]* #2 $n = 22$, Wild-type hermaphrodite $n = 20$, *Ex[inx-18p::fem-3]* #1 $n = 18$, *Ex[inx-18p::fem-3]* #2 $n = 25$ |
| Figures 3C and 3D | Wild-type $n = 24$, *unc-6(e400)* $n = 25$, *unc-40(e271)* $n = 23$, *unc-5(e53)* $n = 25$ |
| Figures 4B and 4C | Wild-type $n = 24$, *unc-40(e271)* $n = 23$, *Ex[inx-18p::unc-40]* $n = 24$, *Ex[inx-18p::unc-40ΔFNIII4-5domain]* #1 $n = 19$, *Ex[inx-18p::unc-40ΔFNIII4-5domain]* #2 $n = 22$ |
| Figure 5E | Wild-type $n = 25$, recCaspase (PCA) #1 $n = 20$, recCaspase (PCA) #2 $n = 20$, recCaspase (PHC) #1 $n = 24$, recCaspase (PHC) #2 $n = 21$, recCaspase (HOA) #1 $n = 21$, recCaspase (HOA) #2 $n = 17$ |
| Figure 6A | Wild-type $n = 20$, *Ex[inx-18p::tra-2ic]* #1 $n = 19$, *Ex[inx-18p::tra-2ic]* #2 $n = 20$ |
| Figure 6B | Wild-type male $n = 20$, *Ex[inx-18p::tra-2ic]* male #1 $n = 20$, *Ex[inx-18p::tra-2ic]* male #2 $n = 20$, Wild-type hermaphrodite $n = 20$, *Ex[inx-18p::tra-2ic]* hermaphrodite #1 $n = 20$, *Ex[inx-18p::tra-2ic]* hermaphrodite #2 $n = 20$ |
| Figure 6C | Wild-type male $n = 33$, *Ex[inx-18p::tra-2ic]* male #1 $n = 35$, *Ex[inx-18p::tra-2ic]* male #2 $n = 35$, Wild-type hermaphrodite $n = 34$, *Ex[inx-18p::tra-2ic]* hermaphrodite #1 $n = 24$, *Ex[inx-18p::tra-2ic]* hermaphrodite #2 $n = 35$ |
| Figure S2B and S2C | Wild-type hermaphrodite L4 $n = 25$, Wild-type hermaphrodite Young adult $n = 15$ |

